# Genomic characterization of *Pseudomonas syringae* pv. *syringae* from Callery pear and the efficiency of associated phages in disease protection

**DOI:** 10.1101/2023.07.11.545637

**Authors:** D. Holtappels, S.A. Abelson, S.C. Nouth, G.E.J. Rickus, J.P. Giller, B. Koskella

## Abstract

*Pseudomonas syringae* is a heterogeneous species complex of plant pathogenic bacteria associated with a wide distribution of plant species. Advances in genomics are revealing the complex evolutionary history of this species complex and the wide array of genetic adaptations underpinning their diverse lifestyles. Here, we genomically characterize two *P. syringae* isolates collected from diseased Callery pears (*Pyrus calleryana*) in Berkeley, California in 2019 and 2022. We also isolated a lytic bacteriophage, which we characterized and evaluated for biocontrol efficiency. Using a multilocus sequence analysis and core genome alignment, we classified the *P. syringae* isolates as members of phylogroup two, related to other strains previously isolated from *Pyrus* and *Prunus*. An analysis of effector proteins demonstrated an evolutionary conservation of effectoromes across isolates classified in PG2, and yet uncovered unique effector profiles for each, including the two newly identified isolates. Whole genome sequencing of the associated phage uncovered a novel phage genus related to Psa phage PHB09 and the *Flaundravirus* genus. Finally, using in planta infection assays, we demonstrate that the phage was equally useful in symptom mitigation of immature pear fruit regardless of the Pss strain tested. Overall, this study demonstrates the diversity of *P. syringae* and their viruses associated with ornamental pear trees, posing spill-over risks to commercial pear trees and the possibility of using phages as biocontrol agents to reduce the impact of disease.

## Introduction

*Pseudomonas syringae* comprises a species complex of plant pathogens infecting a wide variety of naturally occurring and agricultural plant species. Currently, more than 60 pathovars are described based on the plant host and disease symptoms (1). Over the past years, “First disease reports” describing strains within the *Pseudomonas syringae* species complex have been outnumbering any other group of bacterial phytopathogens and are comparable to certain fungi (2). Among them, *Pseudomonas syringae* pv. *syringae* (Pss) is considered one of the most generalist due to its ability to infect upwards of 180 plant species (3), including multiple *Prunus* species, as well as other stone fruits, mango, citrus and pear (4–6). Multiple Pss strains have been found to be pesticide resistant over a wide variety of chemical control measures, especially copper. Furthermore, the occurrence of pesticide resistance appears to be positively correlate with the use of the respective pesticide, making diseases caused by this pathogen particularly challenging to control (3, 7, 8).

In pear specifically, Pss is known to cause a disease referred to as blossom blight. The disease mainly manifests early in the growing season and is characterized by blossom and bud blast, shoot dieback, and stem cankers (9). The disease was first reported in 1914 by Barker and Grove in the West of England (10, 11). In 1934, Wilson described the disease for the first time in Californian pear orchards located in the Sierra Nevada foothills (12). Research on a variety of Pss strains collected from pear were characterized by a low cultivar specificity, and their ability to cause disease in other plant species such as tomato, underlining the generalist nature of this pathogen (3, 13). In line with their generalist nature, pathogenic strains of Pss have been found on grasses in infested orchards, creating a reservoir of bacterial inoculum and demonstrating a potential spill-over effect (14). Similarly, the Callery pear (*Pyrus calleryana*), an ornamental pear species originating from Asia that has been introduced in the United States in the first half of the 20^th^ century, can act as host to a number of pathogens capable of infecting commercial pears. As these pear trees are susceptible to similar diseases as commercial pear cultivars, such as fire and blossom blight, they pose spill-over risks to spread diseases to commercial and local orchards. Indeed, Pss strain FF5, a common laboratory strain, was initially isolated from Callery pear in Oklahoma, being infectious to both these tree and commercial pears (15).

Recent interest in bacteriophages, viruses of bacteria, as a potential biocontrol measure for bacterial infections in crops has shown promise (16). In the case of Pss specifically, several lytic phages (i.e. those that lyse their host cells) have been described in recent years for use as biocontrol. For example, Pinheiro and colleagues, have reported the potential of the commercially available RNA phage φ6 to control *P. syringae* pv. *phaseolicola, P. syringae* pv. *syringae* and *P. syringae* pv. *actinidiae* strains based on its stability and microbiological characteristics (17). Another Pss phage, phage SoKa, showed disease control potential in *ex planta* bioassays to reduce symptom development in citrus fruit (5). In sweet cherry, nine phages were reported to reduce disease up to 50% in cherry leaflets (18). Similarly, phage MR8 reduced bacterial population with about three log units in cherry twigs, demonstrating the potential of the phage in biocontrol (6). To date, no phages have been described for use in control of Pss in pear specifically, despite the increasing prevalence of diseases caused by *P. syringae* (2).

To address this research gap, we isolated Pss strains and an associated lytic phage from Callery pear growing in Berkeley, California. We characterized the Pss strains using multilocus sequencing analysis and whole genome sequencing, placing them within phylogroup 2 of the species complex, and describe their effector repertoire. We similarly characterized the isolated phage using whole genome sequencing, and tested its biocontrol efficacy against strains collected from Callery pear both *in vitro* and using a detached fruit assay.

## Materials and methods

### Bacterium and phage isolations

Diseased leaf samples (black necrotic tissue) were collected from three Callery pear trees (*Pyrus calleryana*) in Berkeley, California in February, March, April, and May 2019 and 2022. The samples (n=24) were weighed and preserved in a phosphate-glycerol buffer (0.03 M NaH_2_PO_4_, 0.07 M Na_2_HPO_4_ 7H_2_O, 1:10 v% glycerol) at -20°C until further processing. Frozen leaf samples were snap thawed at 56°C and homogenized with a MP FastPrep 24 5G homogenizer using ceramic beads, after which a dilution series was plated on King’s Broth (KB) agar plates to isolate single colonies from the culturable fraction of the bacterial community. Plates were incubated for two days at 30°C.

To isolate the viral fraction, the homogenate was filtered (0.45 µm) and spotted (5 µL) on a bacterial lawn of the isolated Pss strains. Samples showing lysis zones on the Pss strain isolated from February 2019 were plated in triplicate with double agar to determine the concentration of phage infecting the respective bacterial host per gram of leaf tissue. Plaques were picked with sterile toothpicks, resuspended in a physiological buffer (10 mM TrisHCl) and plated with double agar. This was repeated for three consecutive rounds. Phages were propagated by infecting an exponentially growing culture of the respective host (at an optical density of 600 nm (OD_600_) of 0.3) in a 1:100 ratio. The coculture was incubated overnight at 30°C and filtered (0.45 µm).

### Genomic characterization of *Pseudomonas syringae* pv. *syringae*

Yellow fluorescent bacterial colonies from samples showing blossom blight symptoms were picked and the bacterial genus was determined by sequencing the 16S rDNA region. Genomic DNA was isolated using the Qiagen Powersoil Pro kit as described by the manufacturer and Illumina sequenced at The SeqCenter (Pittsburgh, PA, US). The reads were trimmed with Trimmomatic (v0.38) (19) and the quality was assessed with FastQC (v0.11.9). Trimmed reads were assembled with Unicycler (v0.5.0) (20) and annotated using Prokka (v1.14.6)(21). A multilocus sequencing analysis (MLSA) was performed based on the concatenated gyrB and rpoD sequences (3) using MEGA11 (v11.0.11)(22). Sequences were aligned with MUSCLE, followed by constructing a neighbor-joining tree with 1,000 bootstraps and visualized with iTol (v6.6)(23).

A core genome analysis was built by extracting *Pseudomonas syringae* genomes from NCBI (December 2022). The quality of the assemblies was evaluated using BUSCO and genomes of a score above 0.95 were selected for downstream analysis. An ANI analysis was performed by means of fastANI (v1.33)(24). The core genome was determined using Ppanggolin (v.1.2.74) (25) and a phylogenetic tree was constructed using FastTree (26) with default parameters. Similarly, the effector analysis was performed using the effector database from Dillon *et al*. (27). In short, a Blastp analysis was conducted on strains closest related to Pss11.5 and Pss16QsV supplemented with strains originating from pear (ff5, B310D, B310D-R, and LMG5084) with a threshold of 50% query coverage and an e-value of 1e-20 to ensure a conservative annotation of putative effector proteins. Additionally, the protein sequences were manually curated to limit the number of false positives in the dataset. An absence presence matrix for all the different effectors was built and analyzed using the JMP Pro 16 software by a default hierarchical clustering method.

### Phage characterization

The lysis of an exponentially growing culture of Pss 16QsV (OD_600_ at 0.1) was followed over time after infecting the culture with different multiplicities of infection (MOI of 0.1 – 1 – 10) in six-fold using an Infinite M nano Tecan plate reader. The adsorption kinetics of the phage on each of two Pss strains were assessed by infecting a culture at exponential growth phase (OD_600_ of 0.3) with an MOI of 0.01. Samples were taken directly after and ten minutes after inoculation and treated with chloroform, to kill bacterial cells and halt any active phage replication. Phage particles were enumerated using a double agar overlay, and the number of free phage (PFUs; plaque forming units) was calculated. The adsorption constant was determined according to Hyman and Abedon (28).

Phage genomic DNA was extracted by means of a phenol/chloroform/isoamyl alcohol extraction and purified using an ethanol precipitation. The genomic DNA was Illumina sequenced (The SeqCenter, Pittsburgh PA) and reads were trimmed as processed as described above. Assembled contigs were automatically annotated using the Phage algorithm of the PATRICBRC server (29) and manually curated. We taxonomically classified the phage using the VipTree server (30) and VIRIDIC (31). Genome maps were generated with EasyFig (32). The genome of phage 16Q is available at NCBI with accession number OR001909 and bacterial genomes are available under NCBI accession numbers SAMN35531629 and SUB13483132.

### Ex planta bioassay for assessing the potential of phage 16Q as a biocontrol agent

Pathogenicity of the isolated Pss strains and biocontrol potential of the phage were assessed using an unripe pear assay. Unripe pears were collected from the University of California Gill Tract Community Farm, and surface sterilized by submerging them in 1 % bleach for five minutes followed by three rinses with sterile mQ water. Afterwards, the pears were washed with 70% ethanol and dried in a laminar flow. Pss strains were grown in liquid KB medium, centrifuged at 4000 g at 4C and resuspended in phosphate buffered saline (PBS). The fruits were pierced with sterile toothpicks and inoculated with 10^8^ CFU/mL (5 µL) of bacterial suspension at the site of injury across three replicates per pear. Negative controls were inoculated with PBS buffer. The efficiency and local adaptation (i.e. whether phages were more infective to strains collected from the same tree in the same year) of the phages was assessed by adding five microliters of phage suspension (10^8^ PFU/mL and 10^9^ PFU/mL) prior to inoculating the bacteria (10^8^ CFU/mL) in triplicate. The pears were incubated for five days at 30°C in sterile plastic containers. The surface of the diseased area was measured, and log transformed for analysis. An analysis of variance (ANOVA) model considering random effects for individual pears was used to determine the effect of phage treatment on disease outcome using the JMP pro 16 software.

## Results

### Ornamental Callery pear trees in Berkeley suffer from blossom blight caused by *Pseudomonas syringae* pv. *syringae* as confirmed by a MLSA and core genome alignment

Diseased samples from Callery pear trees collected from March 2019 and March 2022 showed presence of *Pseudomonas*-like colonies when plated on KB medium that were confirmed to be *Pseudomonads* by sequencing of the 16S region. A multilocus sequencing analysis based on the gyrB and rpoD gene sequence, according to Guttierez-Barnanquero and colleagues, demonstrated that indeed the two strains isolated cluster together with other *Pseudomonas syringae* pv. *syringae* strains within phylogroup 2 (Fig. 1) (1, 3). As such, our analysis confirmed the presence of Pss in Callery pear in the city of Berkeley (CA) and hence the disease blossom blight. Based on this phylogenetic analysis, Pss16QsV (collected in March 2019) and Pss11.5 (collected in March 2022) were most closely related to Pss2676, PssUNP345, PssUNP349 and PssUNP346, all isolated from bean, emphasizing the generalist nature of Pss phylogroup 2. When only considering those strains collected from pear, Pss16QsV and Pss11.5 were most closely related to Pss7D46, Pss8B48, Pss8C43, Pss7B12 and PssEPSMV3. Other isolates collected from pear trees included in this analysis clustered together in a separate subcluster within phylogroup 2, underscoring the diversity of this specific phylogroup.

**Figure 1.**
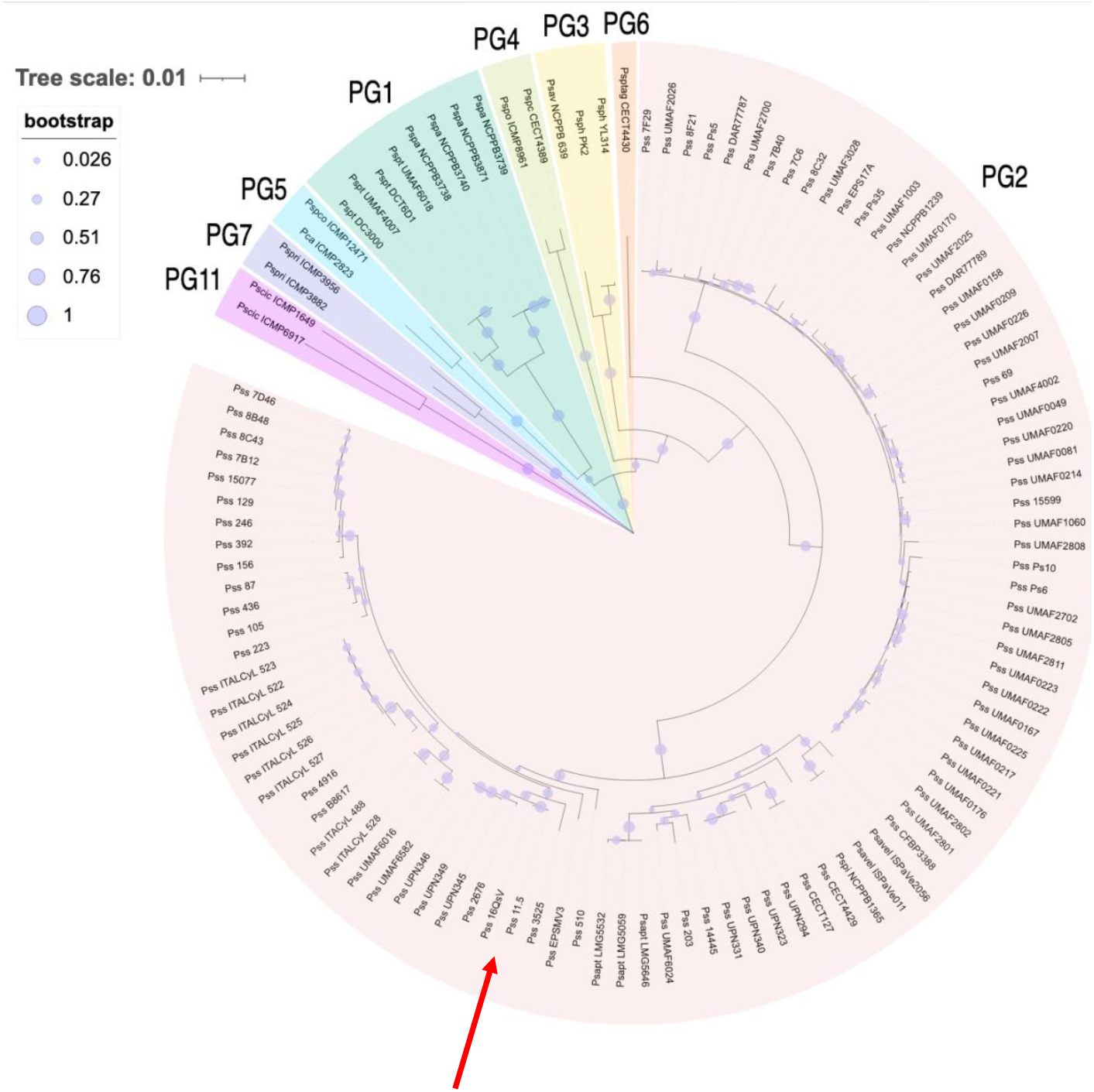
Multilocus sequence analysis of *Pseudomonas syringae* strains according to Gutierrez-Barnanquero et al. including phylogroups 1, 2, 3, 4, 5, 6, 7 and 11 as outgroup. The neighbor-joining tree (1,000 bootstraps) was constructed based on an alignment of partial gyrB and rpoD sequences. The bootstrap values are shown on every node as proportional circles. The evolutionary distance is shown in units of nucleotide substitution per site. The different phylogroups are shown as follows: phylogroup 1 (PG1) in teal, phylogroup 2 (PG2) in red, phylogroup 3 (PG3) in yellow, phylogroup 4 (PG4) in green, phylogroup 5 (PG5) in blue, phylogroup 6 (PG6) in orange, phylogroup 7 (PG7) in purple and phylogroup 11 (PG11) in violet. Pss strains Pss16QsV and Pss11.5 cluster together within PG2 (indicated with red arrow). The phylogenetic tree was generated with MEGA11 and visualized with iTol.

A more in-depth analysis of the genomic diversity of the isolated strains was conducted. To this end, over 600 genomes were extracted from NCBI and analyzed for their completeness using BUSCO. Genomes with a completeness over 95% were used to characterize the pangenome and determine the core genome (n=485). Based on ANI, we identified that our isolates Pss11.5 and Pss16QsV share at least 95% nucleotide identity with other members of PG2, except strain 642 with which our strains share 94% (Supplementary Table 1). According to our pangenome analysis, the pangenome of *P. syringae* consisted of 2,606,355 genes grouped in a total of 73,863 gene families. From these families, 2,637 were persistent gene families (shared between 90 and 100%), 8,470 were shell gene families (shared between 15 and 90% of genomes) and 62,756 were cloud gene families (shared <15% of genomes). Figure 2 shows a phylogenetic tree based on the persistent gene families. Similar to our MLSA, a grouping of the genomes was observed that resembled the different phylogroups as defined for the species of *P. syringae* as reported previously (27). The strains collected in this study clustered together with other strains defined as members of Phylogroup 2, further confirming the results obtained from our MLSA.

**Figure 2.**
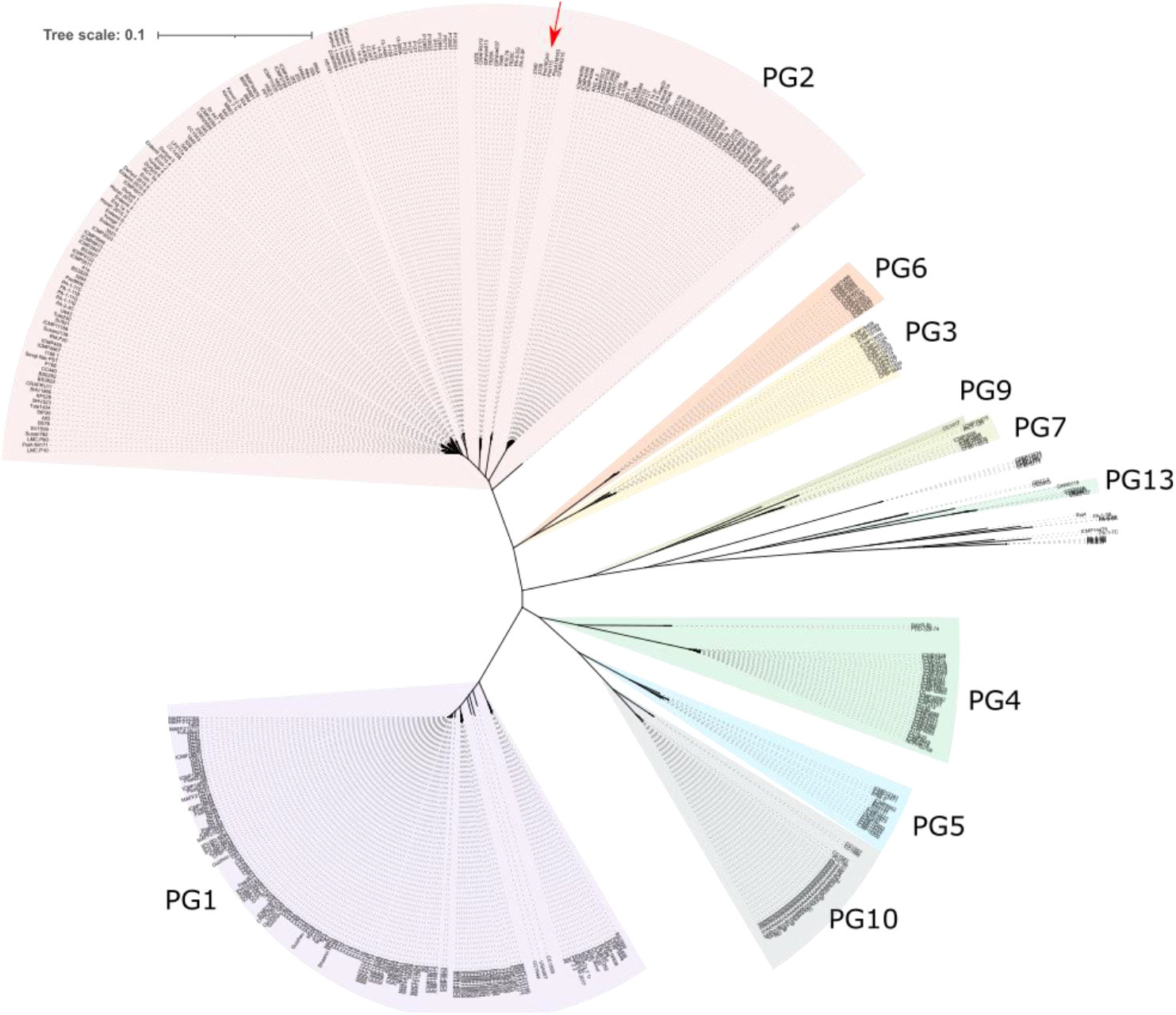
Unrooted phylogenetic tree (n=485 isolates) based on an alignment of the persistent gene families within *P. syringae*. Based on their phylogenetic distance, isolates cluster together according to their phylogroup (purple is phylogroup 1 PG1, pink is phylogroup 2 PG2, orange is phylogroup 6 PG6, yellow is phylogroup 3 PG3, olive green is phylogroup 9 PG9, green is phylogroup 7 PG7, teal is phylogroup 13 PG13, dark teal is phylogroup 4 PG4, blue is phylogroup 5 PG5 and grey is phylogroup 10 PG10). Strains Pss16QsV and Pss11.5 are indicated with a red arrow.

Based on their core genome, Pss16QsV and Pss11.5 were closest related to CFBP4215, PssA1M163, NRS 2339, and NRS 2340 (over 98% ANI). All these strains originated from *Prunus avium* (sweet cherry), except NRS 2340 which was originally isolated from pear.

### Effectoromes are non-random within the phylogeny of *P. syringae*, and Pss16QsV and Pss11.5 encode effector proteins unique for their clade

Effector proteins are considered one of the main virulence factors of *P. syringae*. Hence, we mined the genomes of the closest relatives (n=100) of Pss16QsV and Pss11.5 supplemented with other strains isolated from pear (ff5, B301D, B301D-R, and LMG5084). According to our conservative bioinformatical pipeline, the isolates had on average 18 effector proteins encoded in their genomes with Pss FF5 the lowest amount (n=11) and strain 203 the highest number (n=27). Pss16QsV, Pss11.5, and LMG5084 were annotated to encode 16 effector proteins in their genomes which is higher compared to other strains isolated from pear. Our overall analysis on the effectoromes of PG2 *P. syringae* isolates showed that all strains encoded HopAA1, HopAG1, HopAH1, HopB4, and HopM-ShcM1. Most strains carried the genes for HopI1 and HopB2 with some minor exceptions. More specifically, HopI1 was not predicted in strains FF5, 13-630A, and 14-Gil, while HopB2 was not encoded by ISPaVe037, LP686.1a, and PA-5-3G. Interestingly, our analysis could not detect *avrE1* in Pss FF5, while the gene was present in all other strains. HopAM1 and HopZ6, on the contrary, were only encoded by strains PP1 and HS191, respectively.

We clustered the strains according to the absence and presence of the effector proteins by means of hierarchical clustering (Fig. 3A). Based on this clustering method, seven different groups were distinguished. Comparing the phylogeny of these strains and their effectorome, clade 1 only contained isolates previously classified as PG2 clade 4. Similarly, group 2 and group 3 included primarily isolates from PG2 clade 1, with strain SZ47 as an exception. Group 4, containing isolates Pss11.5 and Pss16QsV, consisted of PG2 clade 2. Group 5 was mainly composed of strains from PG2 clade 4, and group 6 and 7 included isolates from PG2 clade 4 and 1, respectively. As such, our analysis demonstrated a clear relationship between the different clades within PG2 and their effectorome. Indeed, an analysis of similarities (ANOSIM) showed significance between the effectorome of the clades within PG2 (R=0.614, p-value=0.001) suggesting that effectoromes are not random within the different clades of PG2 and that there is an association between phylogeny and the effectorome. Within these groups, however, there were individual genes detected in isolates that are unique to the overall pattern, demonstrating a gain and loss of individual effector genes and hence plasticity in the effectorome.

**Figure 3.**
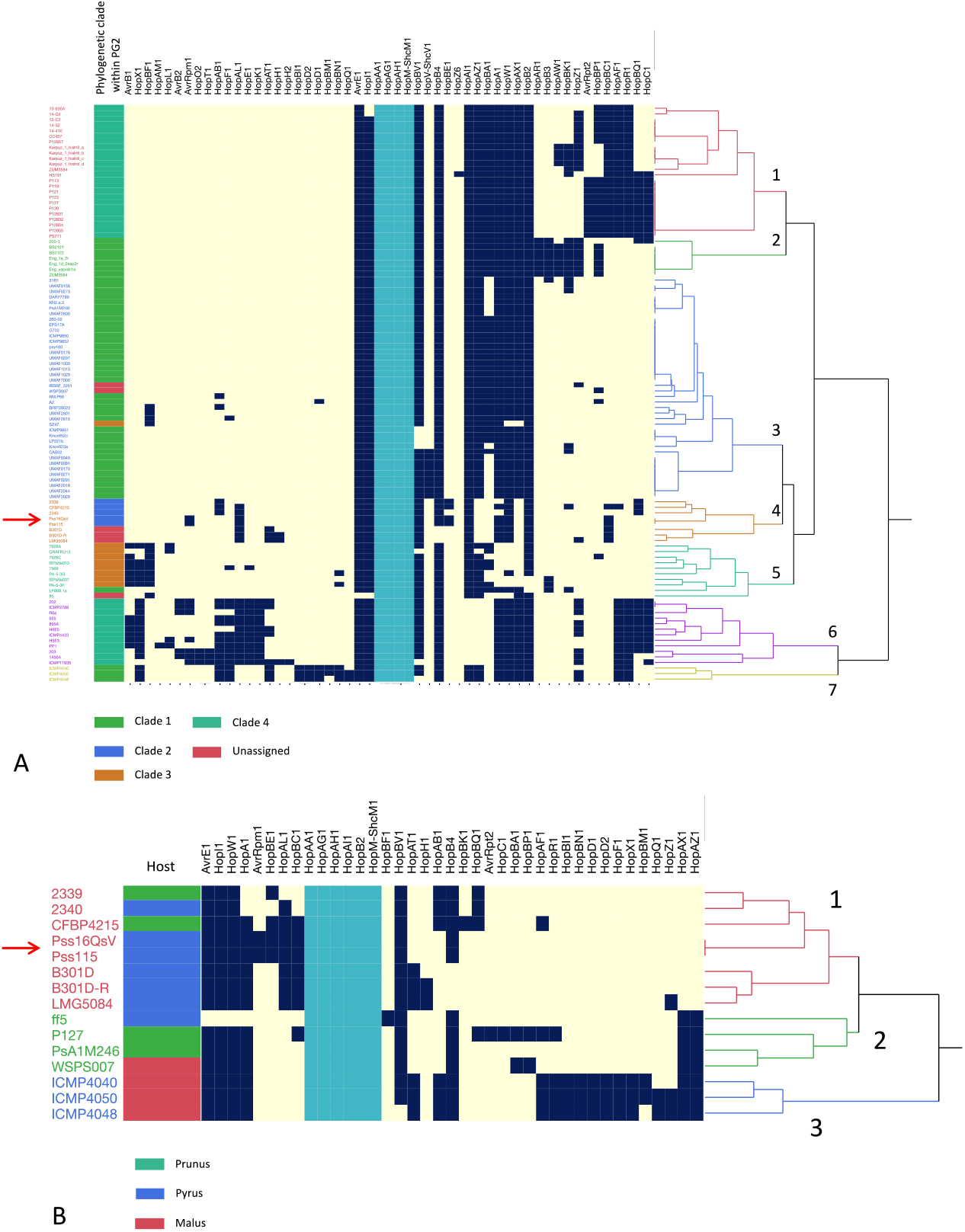
Absence/presence matrix and hierarchical clustering of the screened effector proteins within the different Pss isolates. Yellow indicates if the gene encoding the specific effector protein is not present, dark blue if the gene is encoded in the genome. Strains collected in this research are indicated with a red arrow. Effectors that are encoded by all strains are indicated in light blue. A. The effectorome of a subset of the PG2 strains classified according to our ANI and core genome analysis supplemented with strains specifically isolated from pear (in green clade 1, in blue clade 2, in brown clade 3, in teal clade 4, and in red the unassigned isolates). Based on the A/P of the effectors, seven groups can be distinguished (red, green, blue, orange, teal, and purple). All strains encode HopAA1, HopAG1, HopAH1, HopB4, and HopM-ShcM1. On average, the different strains encode 18 effector proteins. B. In detail the effectorome of Pss isolates isolated from *Rosaceae* trees (in green *Prunus*, in blue *Pyrus*, and in red *Malus*). Within these strains, three different patterns were determined based on the absence and presence of specific effector proteins.

Similar to our phylogenetic analysis, Pss16QsV and Pss11.5 clustered together with CFBP4215, NRS 2339, and NRS 2340, further underlining the close evolutionary relatedness between these strains and the non-randomness of the effectorome. Remarkably, other isolates collected from pear, B301D, B301D-R, and LMG5084 clustered within this group, but not FF5. All strains within group 4 were predicted to encode HopAL1, unique for cluster 3 and cluster 6.

Next, we looked more closely to isolated that were collected from *Rosaceae* species (Fig. 3B). Here, a clear conservation of effector proteins was noticeable within these strains. We can hypothesize that these proteins are essential for *P. syringae*’s association to plants within the *Rosaceae* family. Yet, there was clear strain specificity. Pss11.5 and Pss16QsV, for example, were predicted to encode AvrRpm1 unique for all strains originating from *Rosaceae*. Similarly, strain 2340 was the only pear associated isolate encoding HopAB1. The effector profile of Pss strain FF5 was remarkably distinct from the other strains assessed comprising of HopBF1.

### Novel lytic Pss phage 16Q infects both Pss16QsV and Pss11.5 with similar efficiency and represents a novel phage genus

In addition to the bacteria, we isolated four bacteriophages that were capable of infecting (i.e. forming plaques on a lawn of) strain Pss16QsV from the February (sympatric), March, April, and May 2019 samples. Upon sequencing the phage genomes, we could not identify single nucleotide polymorphisms or other signs of genome plasticity across the phage genomes. As such, we continued our phage characterization with phage 16Q (February 2019), collected in the same sample as Pss16QsV. We evaluated the infection efficiency of the phage in liquid broth on both Pss16QsV (sympatric strain) and Pss11.5 (allopatric strain from a different tree and collected in 2022; Supplementary Fig. 1). This analysis demonstrated that Pss16QsV and Pss11.5 are equally susceptible to phage 16Q in liquid broth. Additionally, we evaluated the adsorption kinetics of phage 16Q on both the sympatric and allopatric strain but found no significant difference in the adsorption of 16Q between the two combinations (Supplementary Figure 2). The adsorption constant for 16Q on both Pss16QsV and Pss11.5 was 7.12e-6 mL/min after ten minutes, suggesting a rather inefficient adsorption of phage 16Q to both hosts.

The genome of 16Q is 94.6 kb with a GC content of 45%. Based on our annotation of the 16Q genome, the phage encoded a panel of 15 tRNAs. Yet, no signs of alternative coding density was detected based on the coding density in alternative translation tables (4, 11, and 15) (33). A phylogenetic analysis of the 16Q genome revealed that this phage is related to the *Pakpunavirus, Otagovirus* and *Flaundravirus* genus (VipTree analysis – Supplementary Figure 3). A more detailed study of the sequence identity between these genera showed that phage 16Q shares over 35% sequence identity with the *Otagovirus* and *Flaundravirus* genera (Fig 4A). Interestingly, all representative members as currently described in the taxonomy database from the International Committee on Taxonomy of Viruses (ICTV – December 2022) infect *P. syringae*, emphasizing their relatively close relatedness and a recent common ancestor. Furthermore, the *Otagovirus* and *Flaundravirus* genera shared about 10% sequence identity with the *Pakpunavirus* genus which is primarily represented by *P. aeruginosa* phages (ICTV – December 2022), again illustrating the relatedness, evolutionary trajectory, and phage-host association. Phage 16Q shared 66.74% identity with its closest neighbor, *P. syringae* pv. *actinidiae* (Psa) phage PHB09. Based on the threshold of sequence identity between phage genera as determined by the ICTV (34), we suggest that phage 16Q represents a new phage genus, closely related to Psa phage PHB09 and the *Flaundravirus* genus.

**Figure 4.**
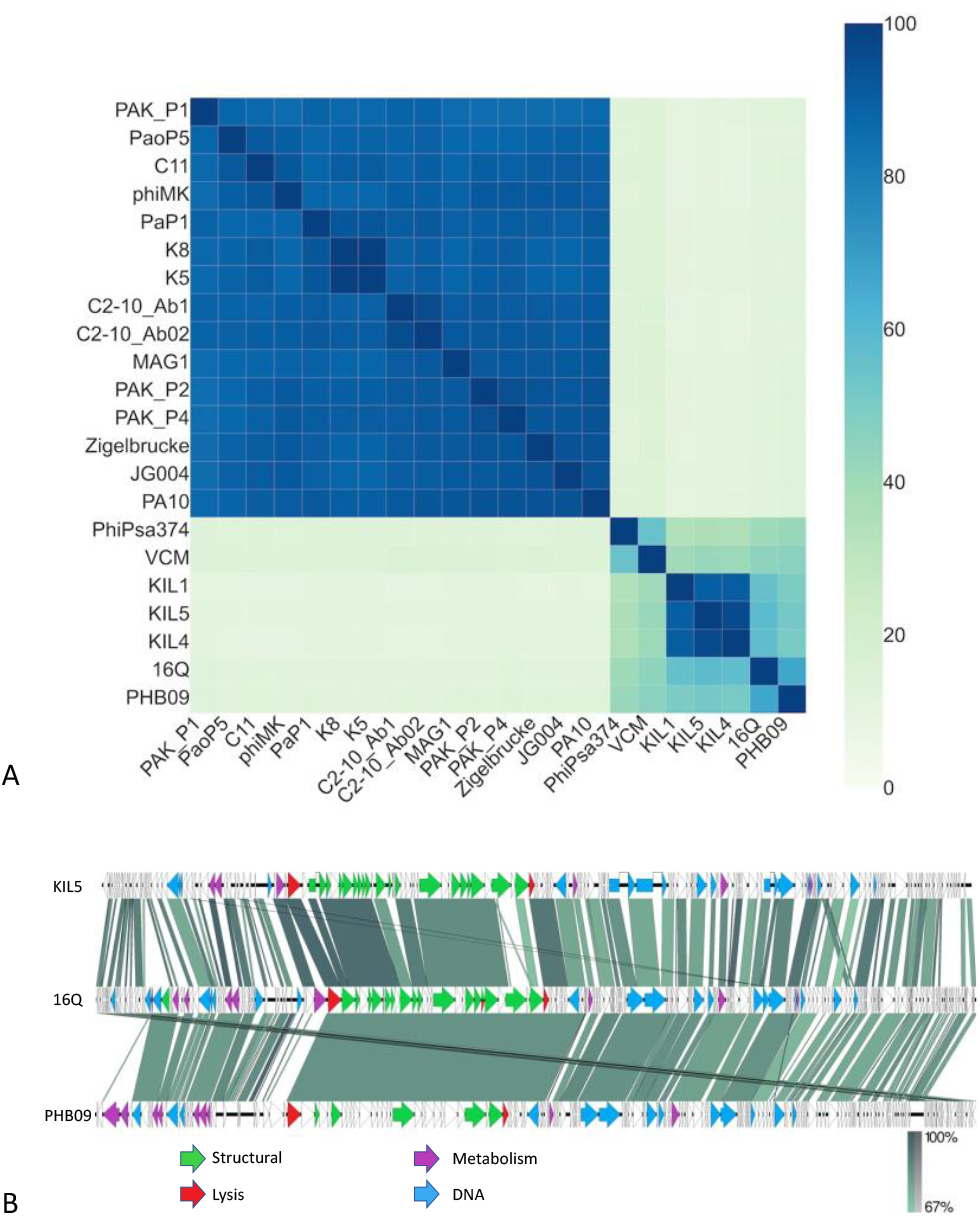
A. Heatmap of a VIRIDIC analysis of phage 16Q along with members of the *Pakpunavirus, Flaundravirus* and *Otagovirus* genera as identified as the closest relatives of phage 16Q based on a VipTree analysis of the 16Q genome. Pss phage 16Q shares 66.74% sequence identity with its closest neighbor *Pseudomonas syringae* pv. *actinidae* phage PHB09 and between 56% and 58% with members of the *Flaundravirus* genus *Pseudomonas syringae* pv. *porri* phages KIL1, KIL4 and KIL5. B. Genome maps of Pspo phage KIL5, Pss phage 16Q and Psa phage PHB09. The phages share a similar genome architecture and homology (blastn) between the different gene clusters. Genes coding for DNA associated proteins are shown in blue, structural proteins in green, lysis associated in red and metabolism associated in violet. Genome orientation and ends of phage 16Q were aligned with Pspo phage KIL5 as these were experimentally determined using primer walking (Rombouts et al. 2016).

This relatedness was further highlighted by comparing the genomes of phage 16Q, *P. syringae* pv. *porri* phage KIL5 and Psa phage PHB09 (Fig 4B). Here, we observed a conservation of genome architecture between the different phages. Similarly, Pspo phage KIL5 and Psa phage PHB09 encoded no genes related to a temperate lifestyle, nor bacterial virulence-associated genes were identified in the genome, suggesting a strictly lytic lifestyle of 16Q and the phage’s potential as a biocontrol agent.

### Phage 16Q shows a significant disease reduction as determined by an *ex planta* bioassay in immature pear fruit

We evaluated the efficiency of phage 16Q in disease protection of both a sympatric and allopatric phage-bacterium combination based on an immature fruit assay. Within our ANOVA model, we found a clear significant increase in the log transformed diameter of the necrotic tissue when the pear fruits were inoculated with Pss (p-value of <0.0001), however, there was no significant difference in the diameter of the necrotic tissue between Pss16QsV and Pss11.5. This suggests that both strains caused similar disease symptoms. When the pears were pretreated with phage, there was a significant reduction of the necrotic tissue developed (p-value of <0.0001). Within our model, the other main effect of the multiplicity in infection (MOI; or ratio of phage to bacterium) was not significant, suggesting that both treatments worked equally well. Similar results were obtained when individual treatments were evaluated against each other using a Tukey-Kramer HSD test correcting for multiple comparisons (connecting letters in Figure 5). Here, there was no significant difference between the two strains, yet, there was a significant effect of the phage treatment. Within the treatments and the allopatric versus sympatric phage-host combinations, there were no significant differences. This suggests that there was no significant effect of local adaptation of phage 16Q in reducing symptoms by two strains.

**Figure 5.**
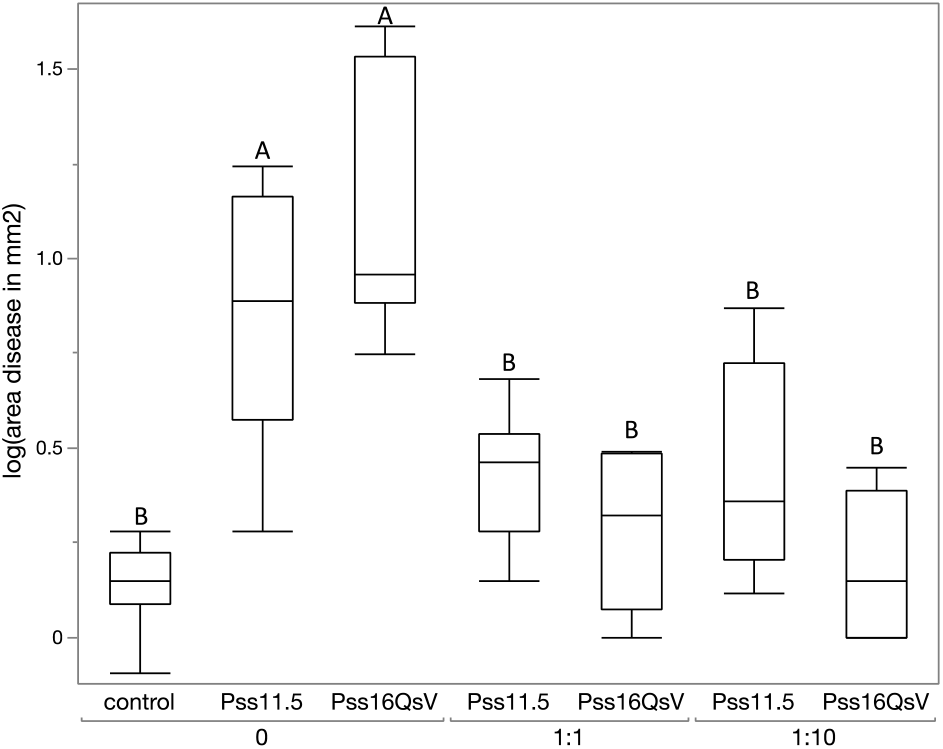
Quantile boxplots of the logarithmic area of necrotic tissue (mm^2^) around the inoculation sites in immature pear fruits (n = 9). Pss16QsV and Pss11.5 were inoculated with a concentration of 10^8^ CFU/mL as well as phage 16Q at a ratio of 1:0 (MOI 0 – no phage), 1:1 (MOI 1) and 1:10 (MOI 10). As negative control, pears were inoculated with PBS. Letters A-B represent different groups of significance (Tukey-Kramer HSD test for multiple comparison, p-value < 0.05).

## Discussion

*Pseudomonas syringae* is a widespread pathogen impacting multiple important crops around the world. In this study, we report and characterize two strains of Pss collected from Callery pear trees exhibiting necrotic symptoms in Berkeley, California. Our immature fruit assays demonstrate that these isolated strains cause disease symptoms in pear, confirming the presence of virulent Pss strains resulting in blossom blight. This was further validated via an MLSA and whole genome sequencing which demonstrate close phylogenetic relatedness with other isolates classified as Pss phylogroup 2. Isolates within this phylogroup are described to infect a wide array of plant species (2, 35). A recent ANI analysis of 99 draft and complete Pss genomes showed that there are three clades within PG2 sharing at least 95% nucleotide identity (36). Looking at the same isolates, we confirm these results. However, including more strains in the analysis reveals the existence of a fourth clade within PG2, indicating that we are only beginning to discover the complex evolutionary history of this phylogroup. On average, we demonstrate that over the whole species complex of *P. syringae* there is a large portion of accessory gene families. This could be explained by the diverse lifestyles and host-associations of isolates within the species complex and the genetic background necessary to support these diverse lifestyles (27, 37).

Essential to the lifestyle of many *P. syringae* isolates are effector proteins. Type III secreted effector proteins, known as the effectorome, are essential in mediating host-microbe interactions as these proteins are translocated to the host cytoplasm where they influence a various cellular processes resulting in host take-over (38). Currently, over 66 families of effector proteins are described for *P. syringae*, demonstrating the diverse strategies this bacterial species has developed over the course of evolution (27). A conservative screening of the absence and presence of effector proteins in the genomes of 494 strains gave clear insights in the number of effector proteins encoded by different phylogroups. As such, the authors found that members of PG2 have a median effector count of 18 effectors per genome, corresponding to our findings (27). The two strains reported in this study encode less than average effector proteins (n=16), yet this number is higher compared to other isolates from pear specifically. Remarkably, we find two effector proteins that are encoded by all strains, including these new isolates. Our analysis demonstrated that HopAH and HopAG are encoded by all PG2 isolates, in line with previous evidence for conservation of these effectors within PG2 (27, 36), suggesting that these effectors are acquired early in PG2 evolution. Indeed, AvrE, HopM, HopAA, and HopB are known to be part of the conserved effector locus (27, 38, 39). We further show that there is more similarity between clades within PG2 than there is among clades, suggesting an evolutionary relatedness of isolates and their effector proteins. Despite this conservation of effectoromes, there is quite some variability in the among the different strains, underlining the importance of horizontal gene transfer of effector genes and the evolutionary arms race between *P. syringae* and its plant hosts (38, 40).

A potential strategy in limiting the spread and impact of Pss is by developing a sustainable biocontrol strategy to contain the disease. To this end, we isolated novel bacteriophages from diseased samples infecting the strains collected from Callery pear. We found one phage (16Q) multiple times (across different trees and collection months), suggesting a widespread distribution of this phage. The genomic analysis of the phage genome revealed that it encodes 15 tRNAs in its genome, making the phage quite independent from its host for translation. This raises the hypothesis that phage 16Q efficiently synchronizes its own transcription and translation which has been hypothesized to be of great importance for efficient host take-over (41) (Holtappels et al. in review). Our phylogenetic analysis of phage 16Q demonstrates a relatedness to other *Pseudomonas* phages, members of the *Pakpunavirus, Otagovirus*, and *Flaudravirus* genera (Figure 3). The former primarily consists of *Pseudomonas aeruginosa* phages while the latter two comprise *Pseudomonas syringae* phages. Interestingly, the overall diversity among phages infecting *P. aeruginosa* is lower than that among phages of *P. syringae*, which corresponds to previous findings of higher diversity among *P. syringae* isolates compared to *P. aeruginosa* strains (42). Remarkably, all seven *P. syringae* phages within the same subcluster as 16Q were reported to be suitable candidates for biocontrol. For example, *P. syringae* pv. *actinidiae (Psa)* phage PhiPsa374 was shown to be stable in storage and did not show any signs of transduction nor a temperate lifestyle. Additionally, this phage had a broad host range, infecting multiple *Psa* strains within the tested collection (43). A more detailed analysis from Warring and colleagues on this phage species demonstrated their effectiveness, both *in vitro* and *in vivo*. They found that the phages adsorbed to lipopolysaccharide residues on the Psa cell surface both *in vitro* and *in planta* and significantly reduced the bacterial load *in planta*. Our immature pear assay demonstrated a similar significant effect of disease mitigation after administering phages, comparable to Pss phage SoKa in citrus (5).

One aspect often overlooked when designing phage therapies is the source of the phage, and whether it comes from the same or different community and/or environment than the intended host. The general concept of phage local adaptation, where phages are found to be more infective to their local host populations than ‘foreign’ ones, has been well described and demonstrated across systems (44–51). In our immature pear fruit, we did not observe a significant difference in the efficiency of the phage to mitigate symptom development in immature fruit, suggesting that phage 16Q is equally well controlling the two strains collected from Callery pear.

## Conclusion

We show that ornamental pear trees function as reservoirs for blossom blight caused by Pss. These strains are characterized by their own unique effector repertoire, arming the bacteria with the virulence factors necessary to infect commercial pear as demonstrated by our immature pear assay. Hence, we conclude that these ornamental pear trees pose a risk of inducing spillovers in urban environments expanding to gardens and commercial orchards, creating a potential biological hazard. However, we also show the promise of phages as biocontrol to reduce such spread, as our newly described 16Q phage reduced symptoms in immature pear fruit across both Pss strains tested. As we continue to uncover the diversity of circulating pathogens and their associated phages, we can move closer to an integrated disease management strategy that takes into account other host reservoirs and alternatives to chemical controls.

## Acknowledgments

The authors would like to thank J.K Sherman for her technical assistance and S.E. Lindow for his contributions to the experimental design. This research was funded by a NSF and USDA/NIFA Career Award (NSF # 1942881, USDA/NIFA # 1024053). DH holds a doctoral fellowship from the “Fonds voor Wetenschappelijk Onderzoek Vlaanderen” (FWO) strategic basic research grant 1S02520N and BK is a Chan Zuckerburg San Francisco BioHub investigator.

---

**Supplementary Figure 1.**
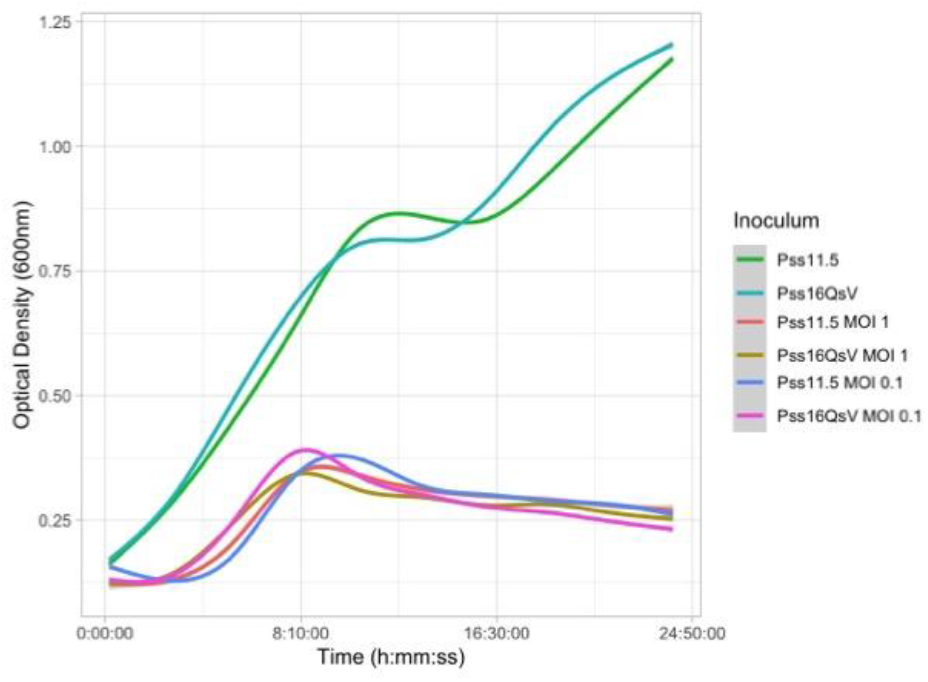
Infection curve (OD_600_ in function of time) of phage 16Q on Pss16QsV and Pss11.5. Phages were added at the start of the measurement at different MOI 0.1 and 1. Around eight hours, bacterial concentrations reached a maximum after which the optical density decreased again.

**Supplementary Figure 2.**
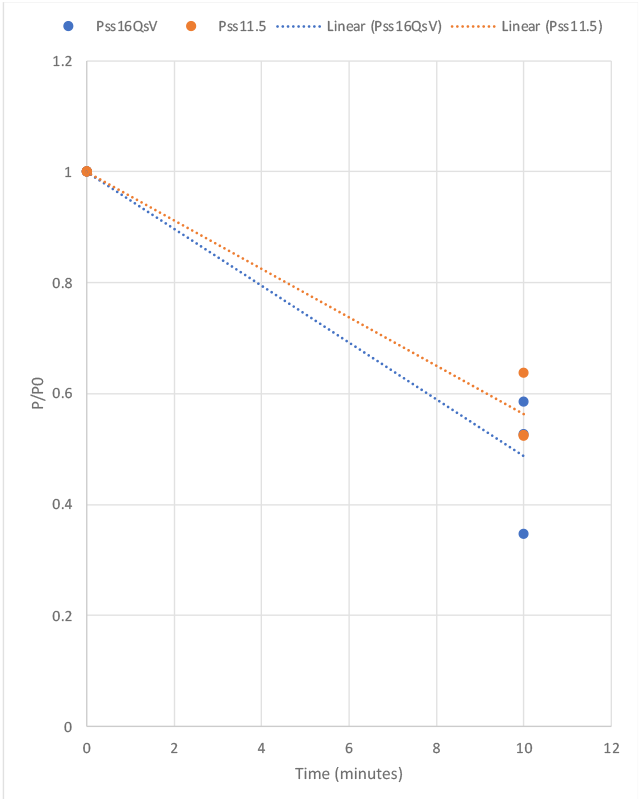
Adsorption curve of phage 16Q on Pss16QsV (Blue) and Pss11.5 (orange) expressed as the ratio of the initial phage concentration and the concentration of free phage after ten minutes in function of time. There was no significant difference in the efficiency of adsorption of phage 16Q on the different strains (p-value >0.05).

**Supplementary Figure 3.**
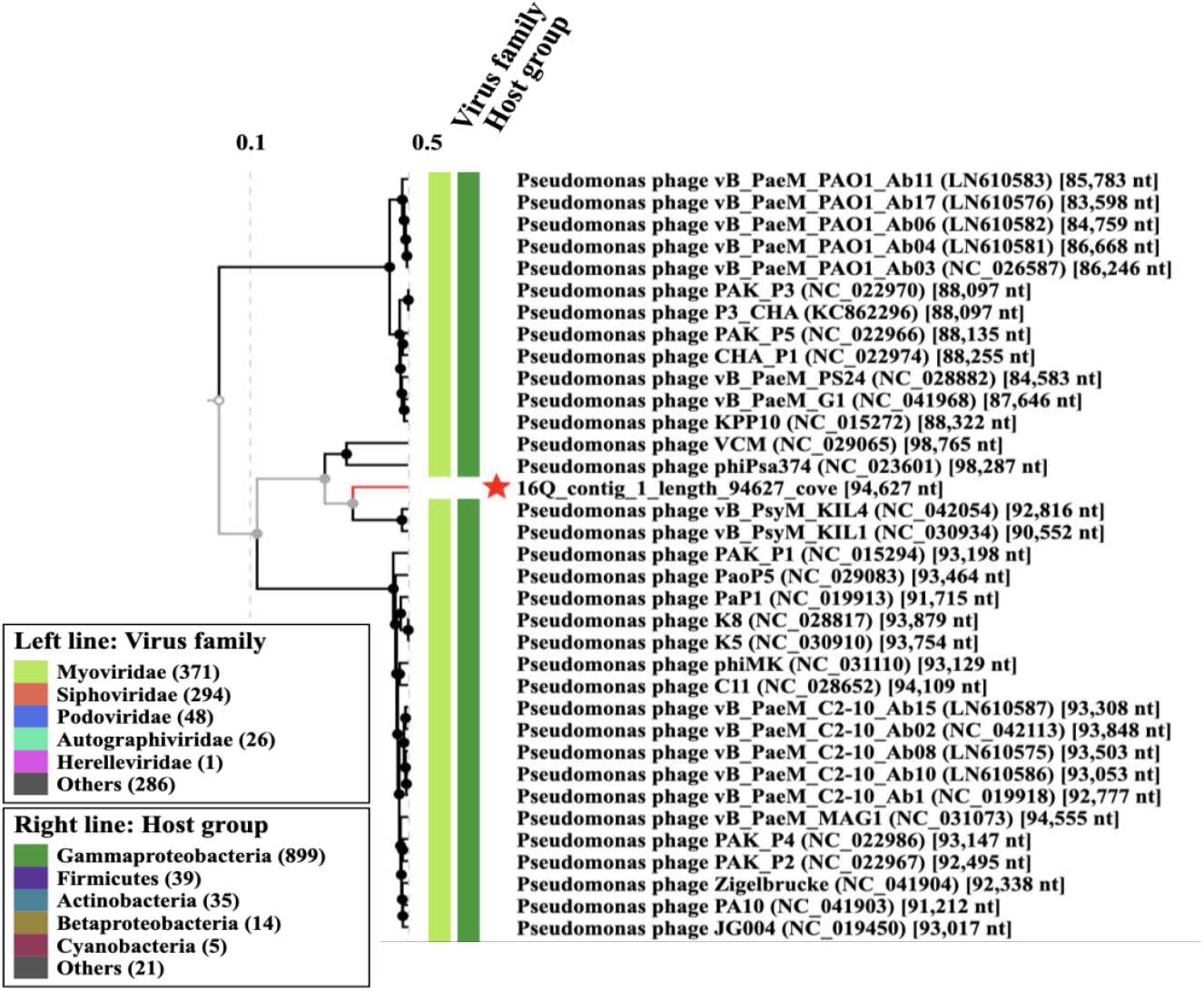
VipTree analysis of phage 16Q. the genome clusters together with members of the Flaundravirus genus as well as phage VCM and phiPsa374.

